# Cyclin G and the Polycomb Repressive Complexes PRC1 and PR-DUB cooperate for developmental stability

**DOI:** 10.1101/144048

**Authors:** Delphine Dardalhon-Cuménal, Jérôme Deraze, Camille A Dupont, Valérie Ribeiro, Anne Coléno-Costes, Juliette Pouch, Stéphane Le Crom, Hélène Thomassin, Vincent Debat, Neel B Randsholt, Frédérique Peronnet

## Abstract

In *Drosophila*, ubiquitous expression of a short Cyclin G isoform generates extreme developmental noise estimated by fluctuating asymmetry (FA), providing a model to tackle developmental stability. This transcriptional cyclin interacts with chromatin regulators of the Enhancer of Trithorax and Polycomb (ETP) and Polycomb families. We investigate here the importance of these interactions in developmental stability. Deregulation of Cyclin G highlights an organ intrinsic control of developmental noise, linked to the ETP-interacting domain, and enhanced by mutations in genes encoding members of the Polycomb Repressive complexes PRC1 and PR-DUB. Deep-sequencing of wing imaginal discs deregulating *CycG* reveals that high developmental noise correlates with up-regulation of genes involved in translation and down-regulation of genes involved in energy production. Most Cyclin G direct transcriptional targets are also direct targets of PRC1, ASX and RNAPolII in the developing wing. Altogether, our results suggest that Cyclin G, PRC1 and PR-DUB cooperate for developmental stability.

## Introduction

Developmental stability has been described as the set of processes that buffer disruption of developmental trajectories for a given genotype within a particular environment (Palmer 1994). In other words, developmental stability compensates the random stochastic variation of processes at play during development. Many mechanisms working from the molecular to the whole organism levels contribute to developmental stability (Nijhout and Davidowitz 2003). For example, chaperones, such as heat-shock proteins, participate in developmental stability by protecting misfolded proteins from denaturation in a large variety of processes (Feder and Hofmann 1999; Queitsch et al. 2002; Rutherford et al. 2007). In *Drosophila*, adjustment of cell growth to cell proliferation is essential to developmental stability by allowing to achieve a consistant organ size (*e.g.* wing size) in spite of variation in cell size or cell number (Debat et al. 2011; Debat and Peronnet 2013).

Developmental noise, the “sum” of the stochastic part of each developmental process, can be observed macroscopically for morphological traits. In bilaterians, quantification of departure from perfect symmetry, the so-called fluctuating asymmetry (FA), is the most commonly used parameter to estimate developmental noise (Van Valen 1962; Palmer and Strobeck 1992). Indeed, the two sides of bilaterally symmetrical traits are influenced by the same genotype and environmental conditions, and differences between them are thus only due to developmental noise. The use of FA as an estimator of developmental noise makes analysis of the mechanistic and genetic bases of developmental stability compatible with custom genetic and molecular approaches of developmental biology.

The genetic bases of developmental stability remain unclear. Thus, its evolutionary role is subject to many speculations (for reviews see Dongen 2006; Leamy and Klingenberg 2015; Debat and Peronnet 2013). Experiments showing the role of *Hsp90* in buffering genetic variation led to the idea that developmental stability could be ensured by specific genes (Rutherford and Lindquist 1998; Milton et al. 2003; Debat et al. 2006; Sangster et al. 2008). On the other hand, both theory and experiments show that complex genetic networks can become intrinsically robust to perturbations, notably through negative and positive feedbacks, suggesting that the topology of gene networks is of paramount importance for developmental stability (Siegal and Bergman 2002). Several authors have further suggested that hubs, *i.e.* the most connected genes in these networks, might be particularly important for developmental stability (Rutherford et al. 2007; Levy and Siegal 2008).

In *Drosophila*, mutants for *dILP8* and *Lgr3* involved in the control of systemic growth, have been reported to display high FA as compared to wild type flies, indicating that these genes are important for developmental stability (Garelli et al. 2012; Colombani et al. 2012, 2015; Vallejo et al. 2015). Two studies have scanned the *Drosophila* genome for regions involved in developmental stability (Breuker et al. 2006; Takahashi et al. 2011). Several deletions increased FA but genes responsible for this effect inside the deletions were not identified. Nevertheless, these studies confirm that the determinism of developmental stability could be polygenic, as suggested by Quantitative Trait Loci analyses in mouse (Leamy et al. 2015 and references therein). Together, these data reinforce the idea that developmental stability depends on gene networks.

We have shown that the gene *Cyclin G* (*CycG*) of *D. melanogaster*, which encodes a protein involved in the cell cycle, is important for developmental stability (Faradji et al. 2011; Debat et al. 2011). Indeed, ubiquitous expression of a short Cyclin G version lacking the C-terminal PEST-rich domain (*CycG*^Δ*P*^) generates a very high FA in several organs, notably in the wing. Interestingly, FA induced by *CycG*Δ*P* expression is associated with loss of correlation between cell size and cell number, suggesting that the noisy process would somehow be connected to cell cycle related cell growth (Debat et al. 2011). Hence, *CycG* deregulation provides a convenient sensitized system to tackle the impact of cell growth on developmental stability.

*CycG* encodes a transcriptional cyclin and interacts with genes of the *Polycomb-group* (*PcG*), *trithorax-group* genes (*trxG*) and *Enhancer of Trithorax and Polycomb* (*ETP*) families (Salvaing et al. 2008). These genes encode evolutionary conserved proteins assembled into large multimeric complexes that bind chromatin. They ensure maintenance of gene expression patterns during development (for a recent review see Geisler and Paro 2015). *PcG* genes are involved in long-term gene repression, whereas *trxG* genes maintain gene activation and counteract *PcG* action. *ETP* genes encode co-factors of both *trxG* and *PcG* genes, and behave alternatively as repressors or activators of target genes (reviewed in Grimaud et al. 2006). More recently, we discovered that *CycG* behaves as an *Enhancer of Polycomb* regarding homeotic gene regulation suggesting that it is involved in the silencing of these genes (Dupont et al. 2015). Importantly, Cyclin G physically interacts with the ETP proteins Additional Sex Comb (ASX) and Corto *via* its N-terminal ETP-interacting domain, and co-localizes with them on polytene chromosomes at many sites. Hence, Cyclin G and these ETPs might share many transcriptional targets and might in particular control cell growth *via* epigenetic regulation of genes involved in growth pathways.

Here, we investigate in depth the role of *CycG* in developmental stability. We first show that localized expression of *CycG*^Δ*P*^ in wing imaginal discs is necessary to induce high FA of adult wings. Furthermore, this organ-autonomous effect increases when the ETP-interacting domain of Cyclin G is removed. We show that several mutations for *PcG* or *ETP* genes, notably those encoding members of the PRC1 and PR-DUB complexes, substantially increase *CycG*-induced FA. Next, we report analysis of the transcriptome of wing imaginal discs expressing *CycG*^Δ*P*^ by RNA-seq and find that transcriptional deregulation of genes involved in translation and energy production correlates with high FA of adult wings. By ChIP-seq, we identify Cyclin G binding sites on the whole genome in wing imaginal discs. Strikingly, we observe a significant overlap with genes also bound by ASX, by the Polycomb Repressive complex PRC1, and by RNAPolII in the same tissue. We identify a sub-network of 222 genes centred on Cyclin G showing simultaneous up-regulation of genes involved in translation and down-regulation of genes involved in mitochondrial activity and metabolism. Taken together, our data suggest that Cyclin G and the Polycomb complexes PRC1 and PR-DUB cooperate in sustaining developmental stability. Coordinated regulation of genes involved in translation and energy production might be important for developmental stability.

## Results

### Expression of CycG^ΔP^ in wing precursors is sufficient to induce high wing FA

We previously reported that expression of *CycG* deleted of the PEST-rich C-terminal domain (amino-acids 541 to 566) (*CycG*^Δ*P*^) under control of ubiquitous drivers (*da-Gal4* or *Actin-Gal4*) generated extremely high FA, notably in wings (Debat et al. 2011) (Fig. 1). The strength of this effect was unprecedented in any system or trait. Expression of *CycG*^Δ*P*^ thus provides a unique tool to investigate developmental stability in depth. To determine whether wing FA was due to local or systemic expression of *CycG*^Δ*P*^, we tested a panel of Gal4 drivers specific for wing imaginal discs or neurons. A brain circuit relaying information for bilateral growth synchronization was recently identified (Vallejo et al. 2015). It notably involves a pair of neurons expressing the dILP8 receptor Lgr3 that connects with the insulin-producing cells (IPCs) and the prothoracicotropic hormone (PTTH) neurons. This circuit was particularly appropriate to test the existence of a remote effect of *CycG*^Δ*P*^ expression in generating high FA in the wing. Expression of *CycG*^Δ*P*^ in this neuronal circuit (using *dilp3-*, *NPF-*, *pdf-, per-, phm-* and *R19B09-Gal4* drivers) did not increase FA of adult wings (Fig. 2 and Sup. Tables 1 and 2). Furthermore, expression of *CycG*^Δ*P*^ in cells of the future wing hinge using the *ts-Gal4* driver did not affect wing FA either. By contrast, expressing *CycG*^Δ*P*^ with 5 different wing pouch drivers (*nub-, omb-, rn-, sd-* and *vg-Gal4*) induced high FA. We thus concluded that *CycG*^Δ*P*^-induced wing FA was due to an intrinsic response of the growing wing tissue.

**Figure 1.**
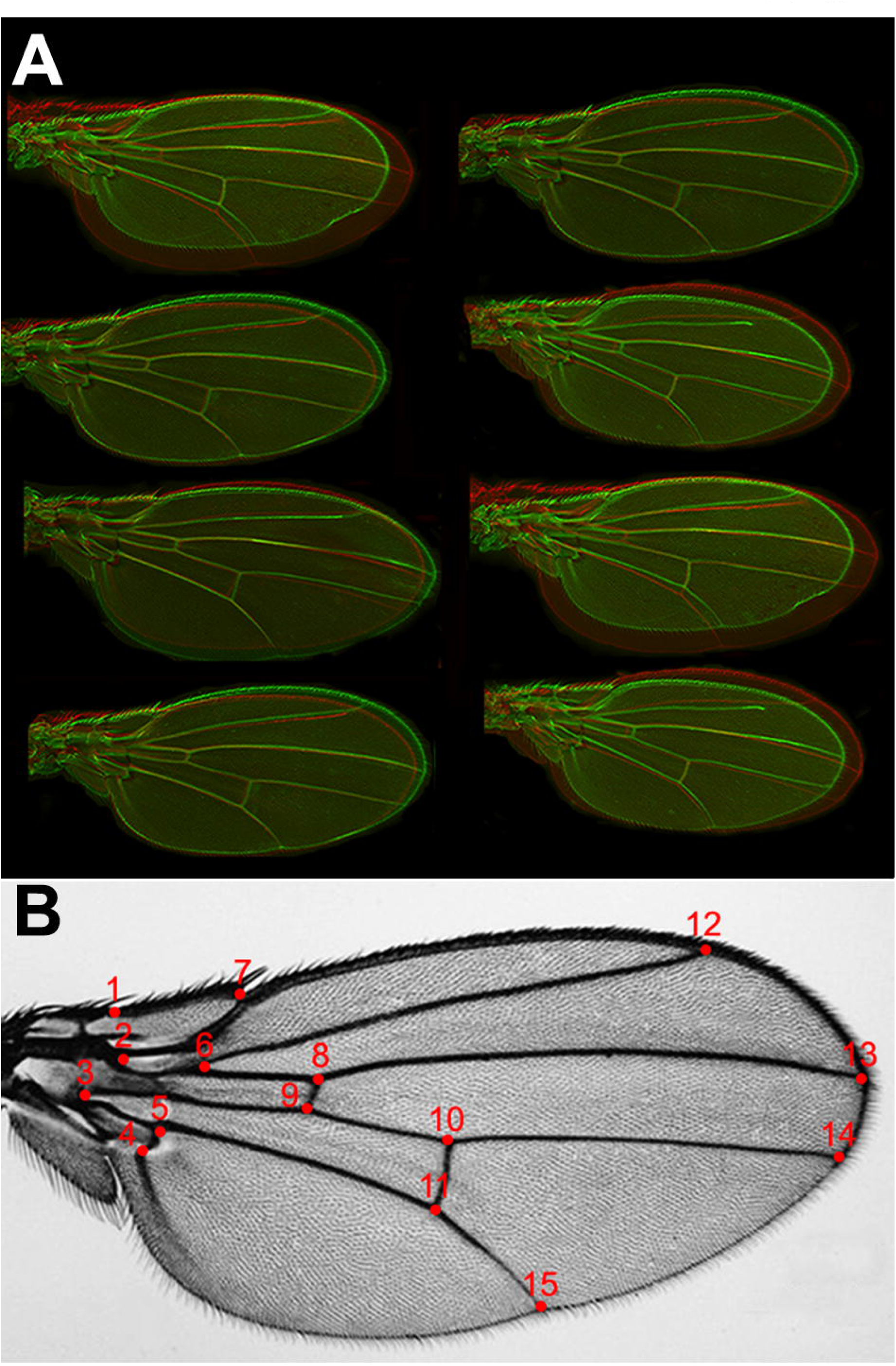
Acquisition of morphometric data. **A –** Superimposition of the left and right wings of a sample of *da-Gal4, UAS-CycG*^Δ*P*^ flies (left wings in red, right wings in green) showing high asymmetry. **B -** Red dots show the 15 landmarks digitized on the wings. The coordinates of these landmarks were obtained from the left and right wings of 30 females randomly sampled from a population. FA was expressed using the FA10 index, *i.e.* the variance of the difference between the left and the right wings in the population, corrected for the measurement error, directional asymmetry and inter-individual variances.

### The Cyclin G ETP interacting domain sustains developmental stability

The 566 amino-acid Cyclin G protein exhibits three structured domains: the ETP-interacting domain (amino-acids 1 to 130) that physically interacts with the ETPs Corto and ASX, a cyclin domain (amino-acids 287 to 360) that presents high similarity with the cyclin domain of vertebrate G-type cyclins, and a PEST-rich domain (amino-acids 541 to 566) (Salvaing et al. 2008; Dupont et al. 2015). To test whether the interaction with ETPs (and thus transcriptional regulation by Cyclin G) could be important to control FA, we generated new transgenic lines enabling to express different versions of the *CycG* cDNA: *CycG*^*FL*^ (encoding the full-length protein), *CycG*^Δ*E*^ (encoding an ETP-interacting domain deleted protein), *CycG*^Δ*P*^ (encoding a PEST domain deleted protein), and *CycG*^Δ*E*Δ*P*^ (encoding an ETP-interacting plus PEST domain deleted protein). In order to express these different cDNAs at the same level and compare the amounts of FA induced, all transgenes were integrated at the same site using the *PhiC31* integrase system. Expression of these transgenic lines was ubiquitously driven by *da-Gal4*. We first confirmed that expression of *CycG*^Δ*P*^ induced very high FA as compared to *+* and *da-Gal4*/+ controls. Furthermore, expression of *CycG*^*FL*^ also significantly increased FA, although to a much lesser extent. Interestingly, expression of either *CycG*Δ*E* or *CycG*Δ*E*Δ*P* significantly increased FA as compared to *CycG*^*FL*^ or *CycG*^Δ*P*^, respectively (Fig. 3 and Sup. Tables 3 and 4). These results show that the ETP interacting domain tends to limit Cyclin G-induced FA and suggest that the interaction between Cyclin G and chromatin regulators sustains developmental stability.

**Figure 2.**
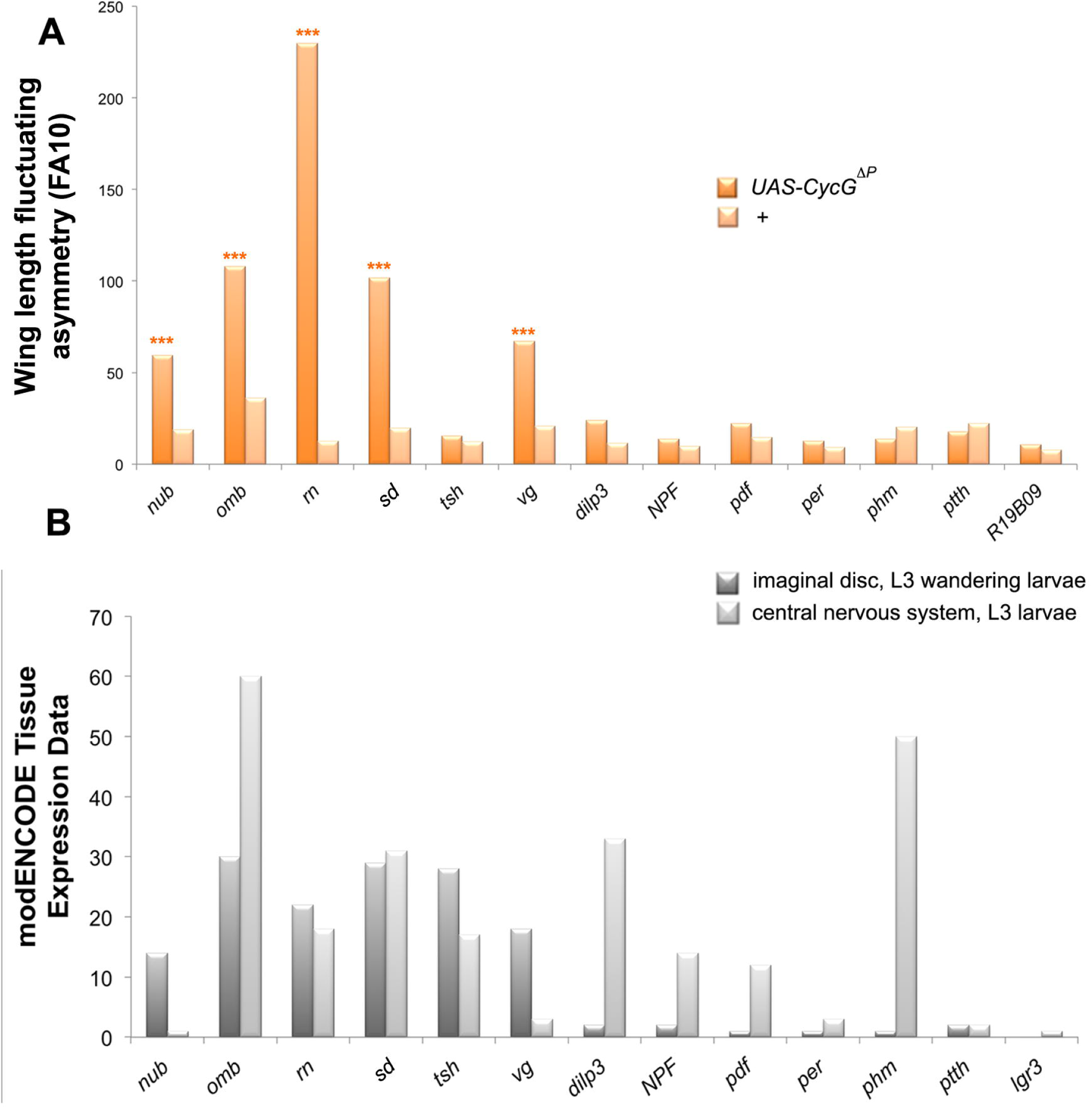
Local deregulation of *CycG* induces high FA. **A –** Wing length FA (FA10) of females bearing a Gal4 driver either associated with *UAS-CycG*^Δ*P*^ (dark orange) or alone (light orange). Wing length was measured as the distance between landmarks 3 and 13. (F-tests, *** p-value<0.001, Sup. Table 1). Source data are provided in Sup. Table 2. **B –** Expression of the driver genes in 3^rd^ instar larva wing imaginal discs (dark grey) and central nervous system (light grey) as indicated in modENCODE (Graveley et al. 2011).

**Figure 3.**
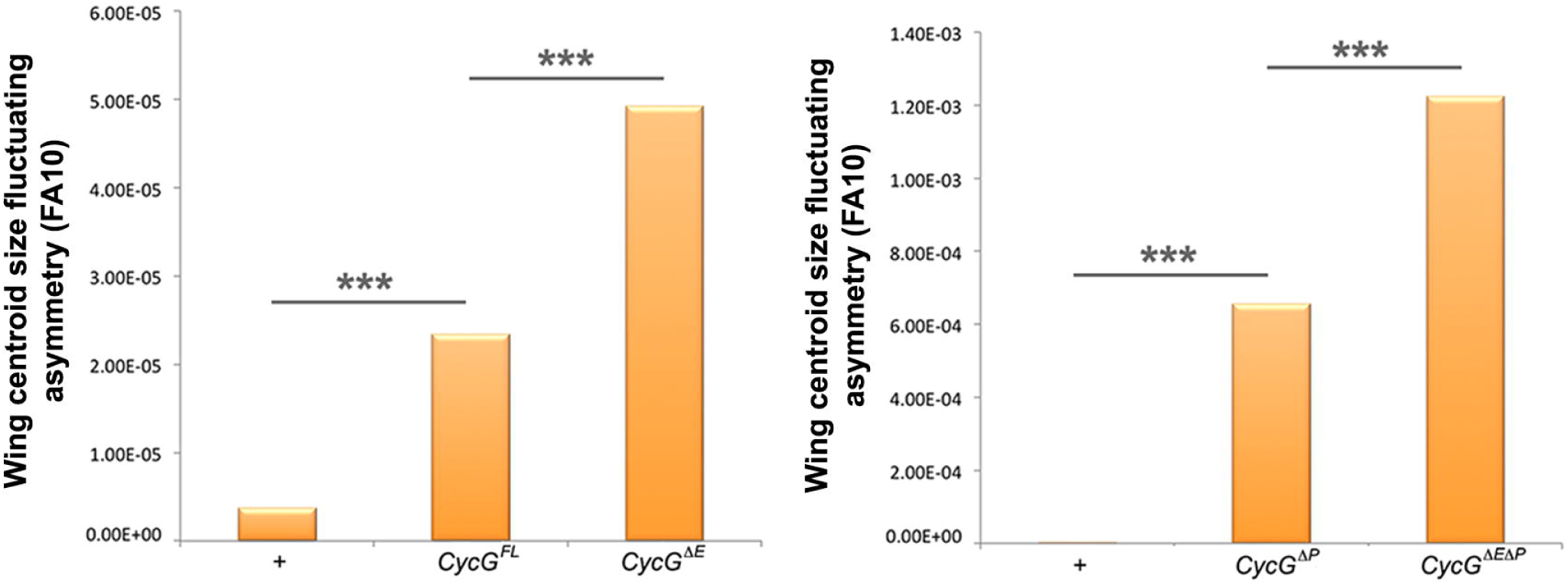
The ETP interacting domain limits *CycG*-induced FA. **A–** Wing centroid size FA (FA10) of females *da-Gal4*/+ (+), *+/UAS-CycG^FL^; da-Gal4/+,* (*CycG*^*FL*^) and *+/UAS-CycG*^Δ*E*^; *da-Gal4,* (*CycG*^Δ*E*^). **B–** Wing centroid size FA (FA10) of females *da-Gal4*/+ (+), *+/UAS-CycG*^Δ*P*^; *da-sGal4/+* (*CycG*^Δ*P*^) and *+/UAS-CycG*^Δ*E*Δ*P*^; *da-Gal4/+* (*CycG*^Δ*E*Δ*P*^). (F-tests, *** p-value<0.001, Sup. Table 3). Source data are provided in Sup. Table 4.

### CycG and PcG or ETP genes interact for developmental stability

We next addressed genetic interactions between *CycG* and *PcG* or *ETP* genes for developmental stability (Table 1). FA of flies heterozygous for *PcG* and *ETP* loss of function alleles was not significantly different from that of control flies. However, when combined with *da-Gal4, UAS-CycG*^*ΔP*^, many of these mutations significantly increased wing FA as compared to *da-Gal4, UAS-CycG*^*ΔP*^ flies (Fig. 4 and Sup. Tables 5 and 6). This was the case for alleles of the PRC1 and PR-DUB encoding genes *Sex comb extra* (*Sce*^*1*^, *Sce*^*33M2*^ and *Sce*^*KO4*^), *calypso* (*caly*^*1*^ and *caly*^*2*^), *Sex comb on midleg* (*Scm*^*D1*^), *Polycomb* (*Pc*^*1*^), and *polyhomeotic* (*ph-p^410^* and *ph-d^401^ph-p^602^*). No modification of *CycGΔP*-induced FA was observed with the *Psc*^*1*^ allele. However, this allele has been described as a complex mutation with loss and gain of function features (Adler et al. 1989). Opposite effects were observed for alleles of the ETPs *Asx* and *corto. Asx^22P4^* increased *CycG*^*ΔP*^ FA whereas *Asx*^*XF23*^ decreased it. *AsxXF*^*23*^ behaves as a null allele but has not been molecularly characterized (Simon et al. 1992), whereas the *Asx*^*22P4*^ allele does not produce any protein and thus reflects the effect of ASX loss (Scheuermann et al. 2010). Similarly, the *corto*^*L1*^ allele increased *CycG*^*ΔP*^-induced FA whereas the *corto*^*420*^ allele had no effect. To characterize these alleles, we combined them with the *Df(3R)6-7* deficiency that uncovers the *corto* locus, amplified the region by PCR and sequenced. The *corto*^*420*^ allele corresponds to a substitution of 14,209 nucleotides starting at position -59 upstream of the *corto* Transcriptional Start Site (TSS) by a 30-nucleotide sequence. Hence, this allele does not produce any truncated protein. By contrast, *corto*^*L1*^ corresponds to a C towards T substitution that introduces a stop codon at position +73 downstream the TSS, generating a 24 amino-acid polypeptide. *corto*^*L1*^ might then behave as a dominant-negative mutation. Lastly, no modification of *CycGΔP*- induced FA was observed for *E(z)^63^* and *esc*^*21*^.

**Figure 4.**
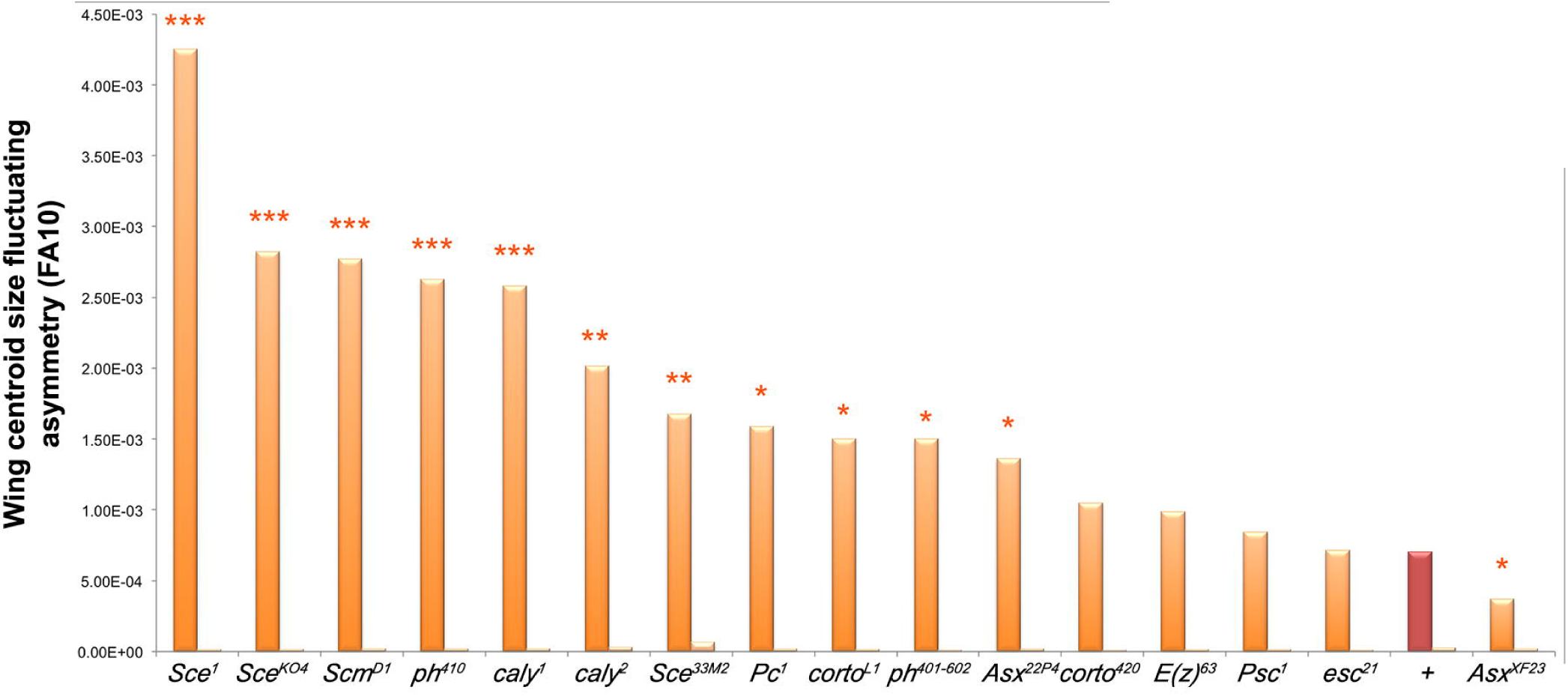
*CycG* interacts with several *PcG* and *ETP* genes for developmental stability. Centroid size FA (FA10) of *ETP* or *PcG* heterozygous mutant females combined with *da-Gal4, UAS-CycG*^*ΔP*^ (dark orange; *da-Gal4, UAS-CycG*^*ΔP*^; *PcG/+* or *da-Gal4, UAS-CycGΔP*; *ETP/+*) and *ETP* or *PcG* heterozygous mutant females combined with *da-Gal4* (light orange; *da-Gal4/+; PcG/+* or *da-Gal4*; *ETP/+*). In brown (*+*), centroid size FA of *da-Gal4, UAS-CycG*^*ΔP*^*/+* and *da-Gal4/+* females. (F-tests, *p-value<0.05; ** p-value<0.01; *** p-value<0.001, Sup. Table 5). Source data are provided in Sup. Table 6.

**Table 1.**
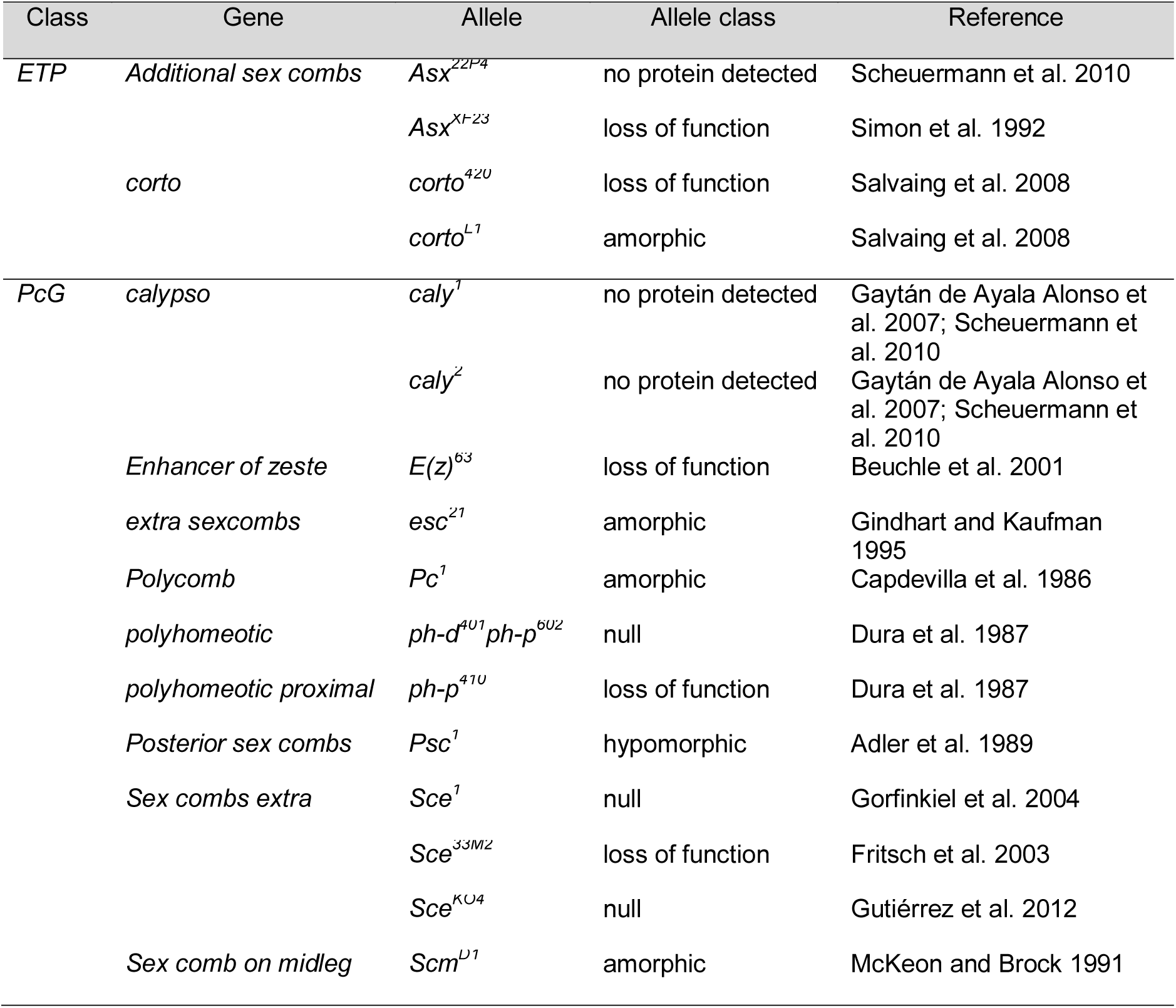
Polycomb and Enhancer of Polycomb and Trithorax alleles used in this study.

Interestingly, *Asx* and *caly* encode proteins of the PR-DUB complex whereas *Pc*, *ph*, *Sce* and *Scm* encode proteins of PRC1, and *E(z)* and *esc* encode proteins of PRC2. Taken together, these results indicate that Cyclin G interacts with the Polycomb complexes PRC1 and PR-DUB, but not with PRC2, for developmental stability.

### *Expression of CycG*^*ΔP*^ *or CycG*^*ΔEΔP*^ *does not modify global H2AK118ub*

Cyclin G binds polytene chromosomes at many sites and co-localizes extensively with PH and ASX suggesting a potential interaction with PRC1 and PR-DUB on chromatin (Salvaing et al. 2008; Dupont et al. 2015). *Sce* and *caly* encode antagonistic enzymes of the PRC1 and PR-DUB complexes, respectively. SCE, aka dRing, ubiquitinates histone H2A on lysine 118 (H2AK118ub) whereas Calypso, aka dBap1, deubiquitinates the same H2A residue (Scheuermann et al. 2010, 2012). To investigate whether Cyclin G was related to these ubiquitin ligase/deubiquitinase activities, we immunostained polytene chromosomes from *w*^*1118*^ larvae with anti-Cyclin G and anti-human H2AK119ub antibodies (homologous to *Drosophila* H2AK118ub) (Lee et al. 2015; Pengelly et al. 2015). Cyclin G and H2AK118ub co-localized extensively on chromosome arms suggesting that Cyclin G transcriptional activity might somehow be connected to this histone mark (Fig. 5A). However, when either *CycG*^*ΔP*^ or *CycG*^*ΔEΔP*^ was expressed in the posterior compartment of wing imaginal discs using the *en-Gal4* driver, the global amount of H2AK118ub was not markedly modified (Fig. 5B, Fig. 5C). We thus concluded that high FA was not related to a global perturbation of H2AK118 ubiquitination.

**Figure 5.**
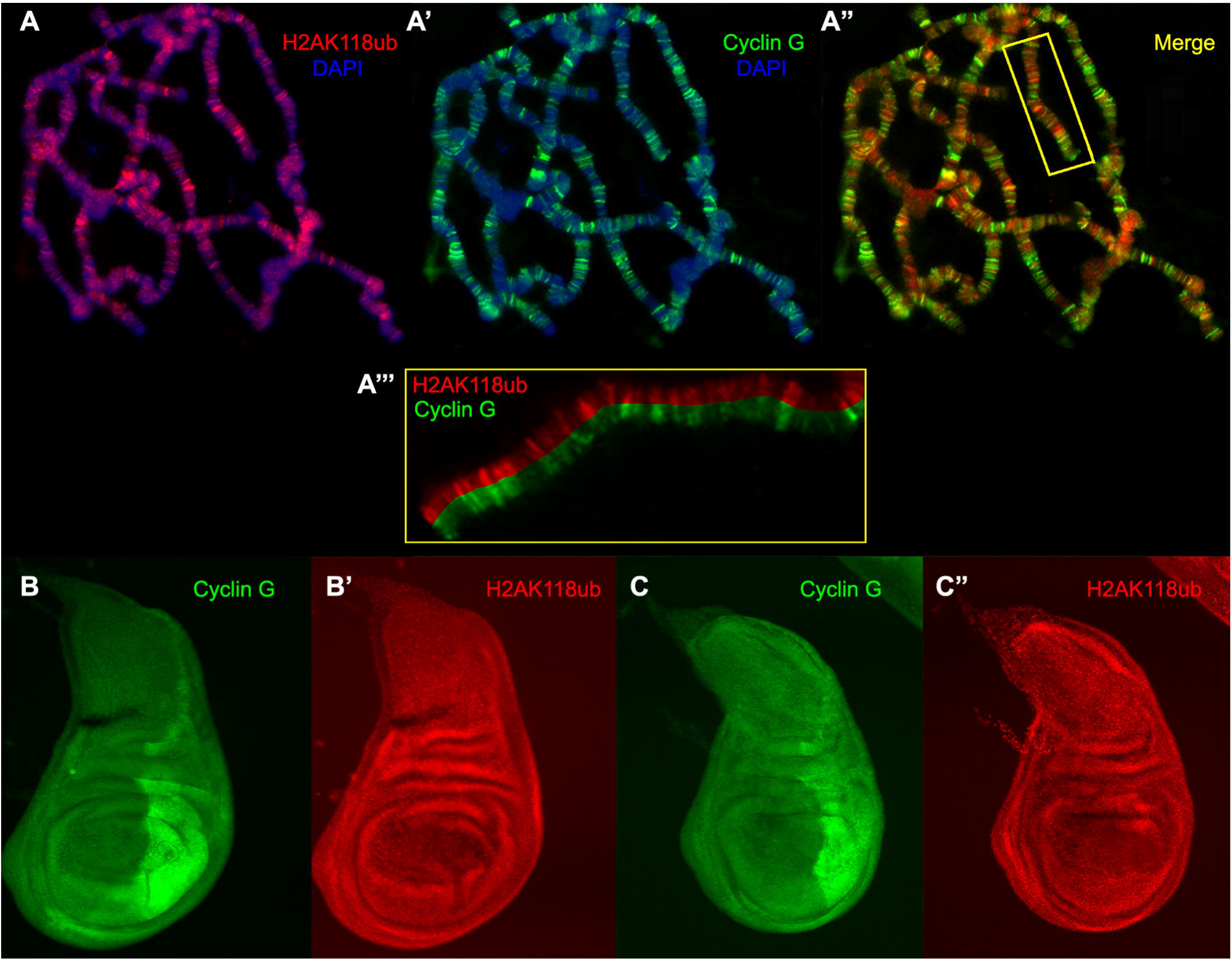
Cyclin G co-localizes with H2AK118ub at many sites on polytene chromosomes but overexpression of *CycG* does not modify global H2AK118ub. **A, A’, A” –** Immunostaining of polytene chromosomes from *w*^*1118*^ third instar larvae. H2AK118ub (red), Cyclin G (green), DAPI (blue). **A’’’ –** Close-up of the box showed in A”. **B, B’ –** Wing imaginal discs of 3^rd^ instar larvae expressing *CycG*^*ΔP*^ in the posterior compartment under control of the *en-Gal4* driver, stained with anti-Cyclin G (green) and anti-H2AK118ub (red). **C, C’ –** Wing imaginal discs of 3^rd^ instar larvae expressing *CycG*^*ΔEΔP*^ in the posterior compartment under control of the *en-Gal4* driver, stained with anti-Cyclin G (green) and anti-H2AK118ub (red).

### Cyclin G controls the expression of genes involved in translation and energy production

Cyclin G controls transcription of the homeotic gene *Abdominal-B* and more specifically behaves as an *Enhancer of PcG* gene for the regulation of homeotic gene expression (Salvaing et al. 2008; Dupont et al. 2015). However, the high number of Cyclin G binding sites on polytene chromosomes suggests that it has many other transcriptional targets. We thus hypothesized that *CycG*^*ΔP*^-induced FA might be related to the deregulation of Cyclin G transcriptional targets. To further address the role of Cyclin G in transcriptional regulation, we deep-sequenced transcripts from *da-Gal4, UAS-CycGΔP/+* and *da-Gal4/+* wing imaginal discs. Sequence reads were aligned with the *D. melanogaster* genome to generate global gene expression profiles. With an adjusted p-value threshold of 0.05, we retrieved 530 genes whose expression was significantly different between *da-Gal4, UAS-CycG*^*ΔP*^*/+* and the *da-Gal4/+* control (Sup. Table 7). Surprisingly, expression of *CycG* was only weakly induced in *da-Gal4, UAS-CycGΔP/+* imaginal discs (1.3 fold). To test the hypothesis that Cyclin G could, directly or not, induce its own repression, we designed primers in the 3’UTR. Indeed, expression of endogenous *CycG* was significantly decreased when *CycG*^*ΔP*^ was expressed (Fig. 6A and Sup. Table 8). Among the 530 genes deregulated in *da-Gal4, UAS-CycG*^*ΔP*^*/+* imaginal discs, 216 were up-regulated and 314 down-regulated. Up-regulated genes were enriched in the Gene Ontology categories *cytoplasmic translation* and *translational initiation* whereas down-regulated genes were enriched in the category *mitochondrial respiratory chain complex* (Fig. 6B and Sup. Table 9). By RT-qPCR, we verified that several ribosomal protein genes (*RpL15, RpL7* and *Rack1*) were over-expressed in *da-Gal4, UAS-CycG^ΔP/^+* imaginal discs (Fig. 6C and Sup. Table 10). In conclusion, *CycG^ΔP^*-induced FA correlates with activation of genes involved in translation and repression of genes involved in energy production.

**Figure 6.**
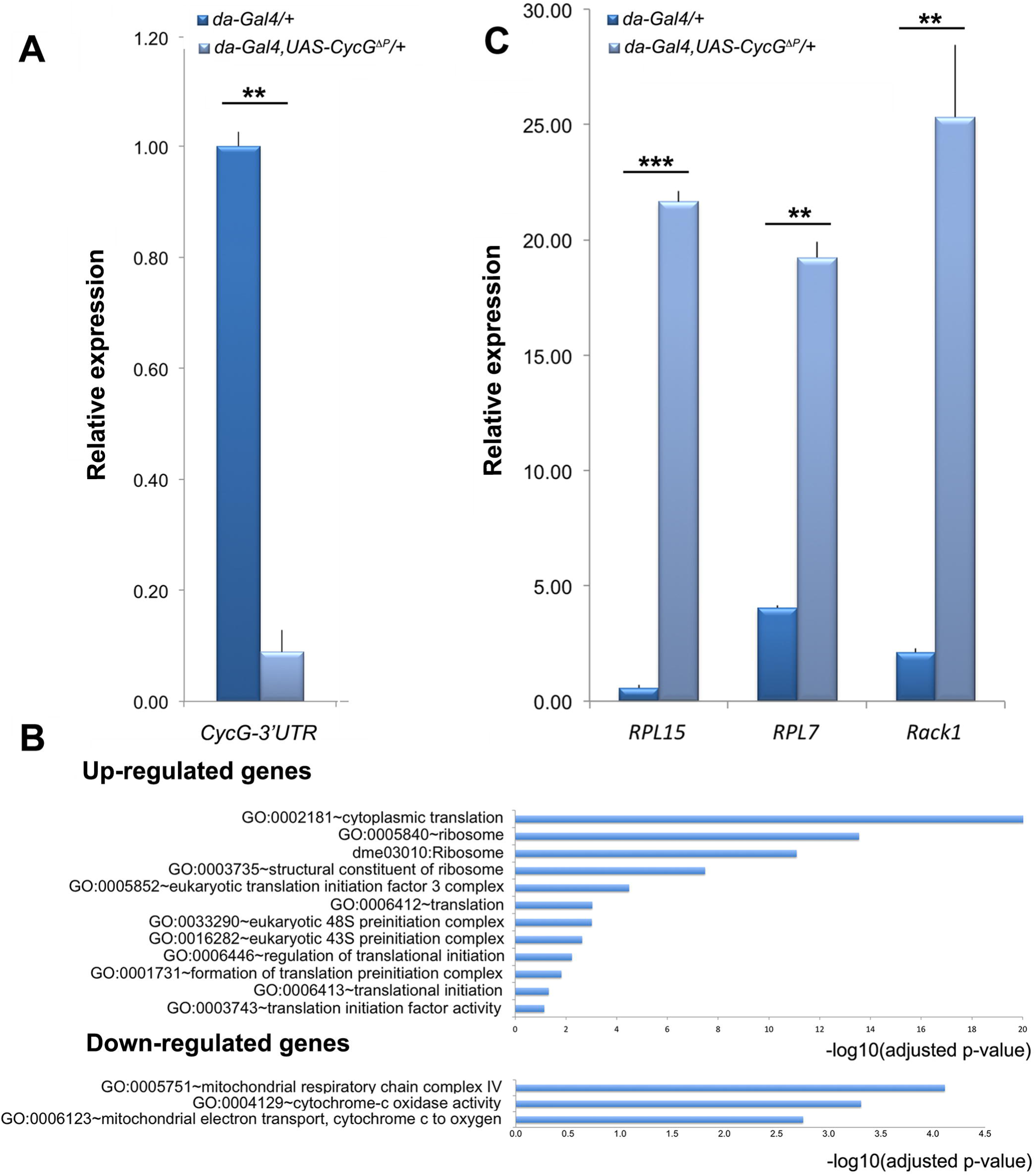
Genes deregulated in wing imaginal discs expressing *CycG*^*ΔP*^. **A –** RT-qPCR analysis of endogenous *CycG* expression in *da-Gal4,UAS-CycG*^*ΔP*^*/+* and *da-Gal4/+* wing imaginal discs. Expression of *CycG* was normalized on the geometric mean of *Lam* and *rin* (Sup. Table 8). t-tests, ** p-value<0.01. Error bars correspond to standard deviations. **B –** Ontology of up-regulated and down-regulated genes in *da-Gal4, UAS-CycG*^*ΔP*^*/+ vs da-Gal4/+* wing imaginal discs. Gene ontology analysis was performed with DAVID (Sup. Table 9). **C –** RT-qPCR analysis of *RPL15*, *RPL7* and *Rack1* expression in *da-Gal4, UAS-CycG*^*ΔP*^*/+* and *da-Gal4/+* wing imaginal discs. Expression of *RPL15*, *RPL7* and *Rack1* were normalized on the geometric mean of *Lam* and *rin* ((Sup. Table 10). t-tests, ** p-value<0.01. Error bars correspond to standard deviations. t-tests, ** p-value<0.01; *** p-value<0.001.

### Cyclin G-bound the TSS of genes involved in translation and protein phosphorylation

To determine the direct transcriptional targets of Cyclin G, we analysed by ChIP-seq the genome-wide binding sites of Cyclin G in *+/UAS-CycG*^*ΔP*^; *da-Gal4/+* imaginal discs. 3363 significant peaks were identified (IDR < 0.05). Among these peaks, 889 were located at TSS of genes whereas the others did not correspond to annotated features (Fig. 7A and 7B and Sup. Table 11). ChIP-qPCR analysis of Cyclin G binding on some of these genes (*RPL7, RPL5, Rack1, CycG*) confirmed that Cyclin peaked on TSS and decreased on gene body (Fig. 7C and Sup. Table 12). As endogenous *CycG* was down-regulated when *CycG*^*ΔP*^ was expressed, we concluded that Cyclin G represses its own promoter.

**Figure 7.**
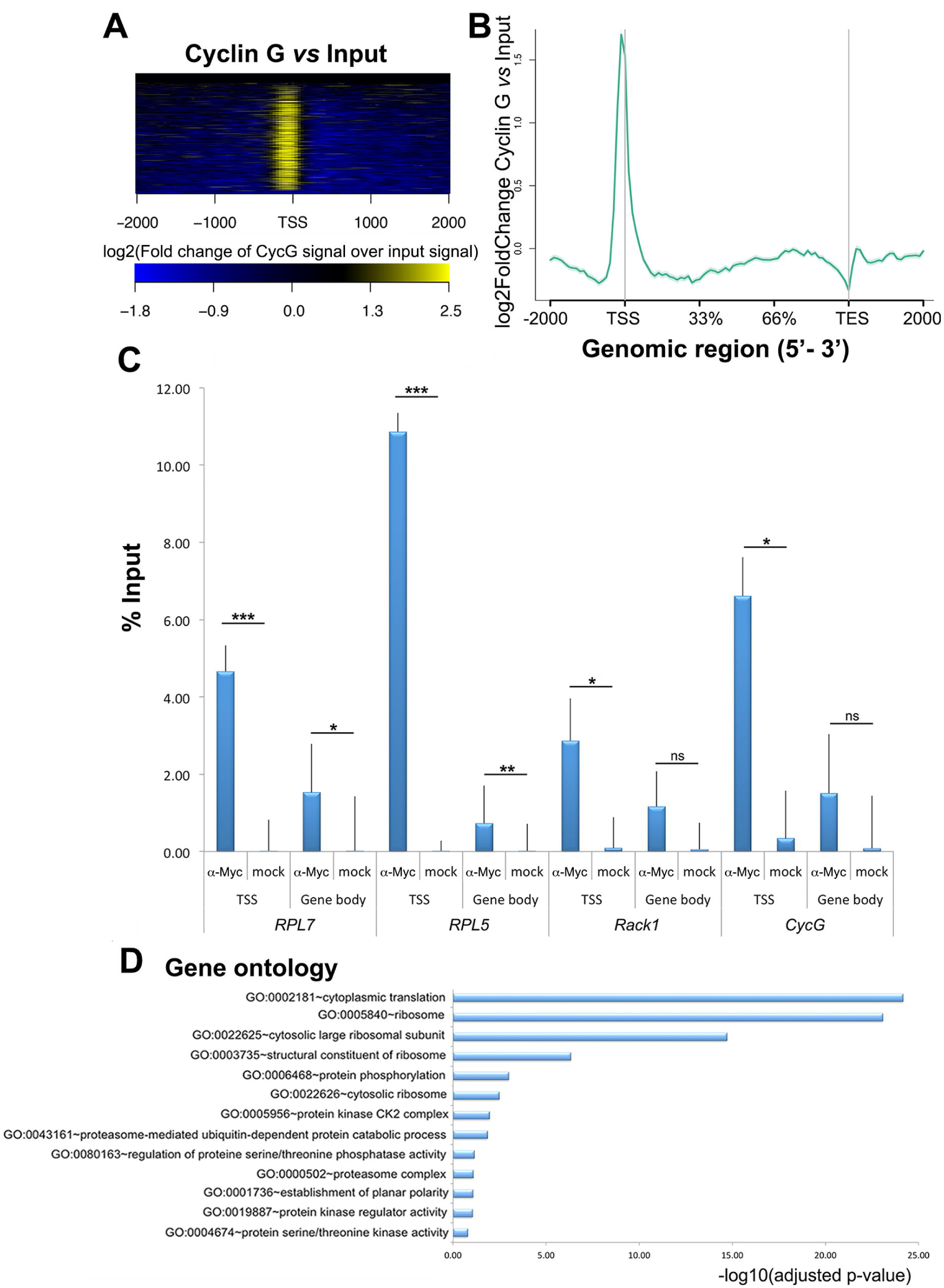
Identification of Cyclin G genome-wide binding sites in wing imaginal discs. **A –** Heat-map showing the enrichment of Cyclin G over the Input signal on the TSS of 889 genes. TSS: Transcriptional Start Site; TES: Transcriptional End Site. **B –** Average profile of Cyclin G signal over these genes shown as an aggregation plot. The standard error is represented as a shaded area over the curve. **C –** ChIP-qPCR analysis of *RPL7, RPL5, Rack1* and *CycG*. IPs were performed either with Myc antibody (α-Myc) to reveal the presence of Cyclin G, or with rabbit IgG as negative control (mock). qPCR were performed using oligonucleotide primers located either at the TSS or in the gene body as indicated (Sup. Table S15). Error bars represent the coefficient of variation (CV) (Sup. Table S12). **D –** Ontology of the 889 genes. Gene ontology analysis was performed with DAVID.

The 889 Cyclin G-bound genes were enriched in GO categories *cytoplasmic translation* and *protein phosphorylation* (Fig. 7D). Comparison of the 530 genes deregulated in imaginal discs expressing *CycG*^*ΔP*^ with the 889 genes presenting a peak at the TSS showed that only 62 genes were both deregulated (39 up- and 23 down-regulated) and bound by Cyclin G (Sup. Table 13). Strikingly, the 39 up-regulated genes were significantly enriched in the GO category *translation* (GO:0002181∼cytoplasmic translation, 14 genes, enrichment score: 11.84, adjusted p-value 2.07E-16).

### Cyclin G-bound genes are enriched in PRC1 and ASX

Using published datasets, we analysed the correlation between regions bound by Cyclin G in +/*UAS-CycGΔP; da-Gal/+* imaginal discs and those enriched in PRC1, or PR-DUB components, RNAPolII, or H3K27me3 in wild type wing imaginal discs (Sup. Table 14). Cyclin G-bound regions were significantly exclusive from H3K27me3, corroborating previous results (Dupont et al. 2015). The same comparisons were performed gene-wise and gave the same results. Importantly, 80% of Cyclin G-bound genes were also bound by RNAPolII (Fig. 8). Considering RNAPolII as a proxy for transcriptional activity, we concluded that Cyclin G-bound genes were located in open chromatin and were either paused or transcribed. However, Cyclin G-bound genes were also significantly enriched in PRC1 target genes. Given that PRC1 has the ability to block transcriptional initiation (Dellino et al. 2004), this indicates that Cyclin G-bound genes were most probably paused. Cyclin G also shared many target genes with ASX but, though ASX and Calypso belong to the PR-DUB complex, Cyclin G did not share binding sites with Calypso. This indicates either that the interaction between Cyclin G and ASX destabilizes the PR-DUB complex or that it takes place outside PR-DUB.

**Figure 8.**
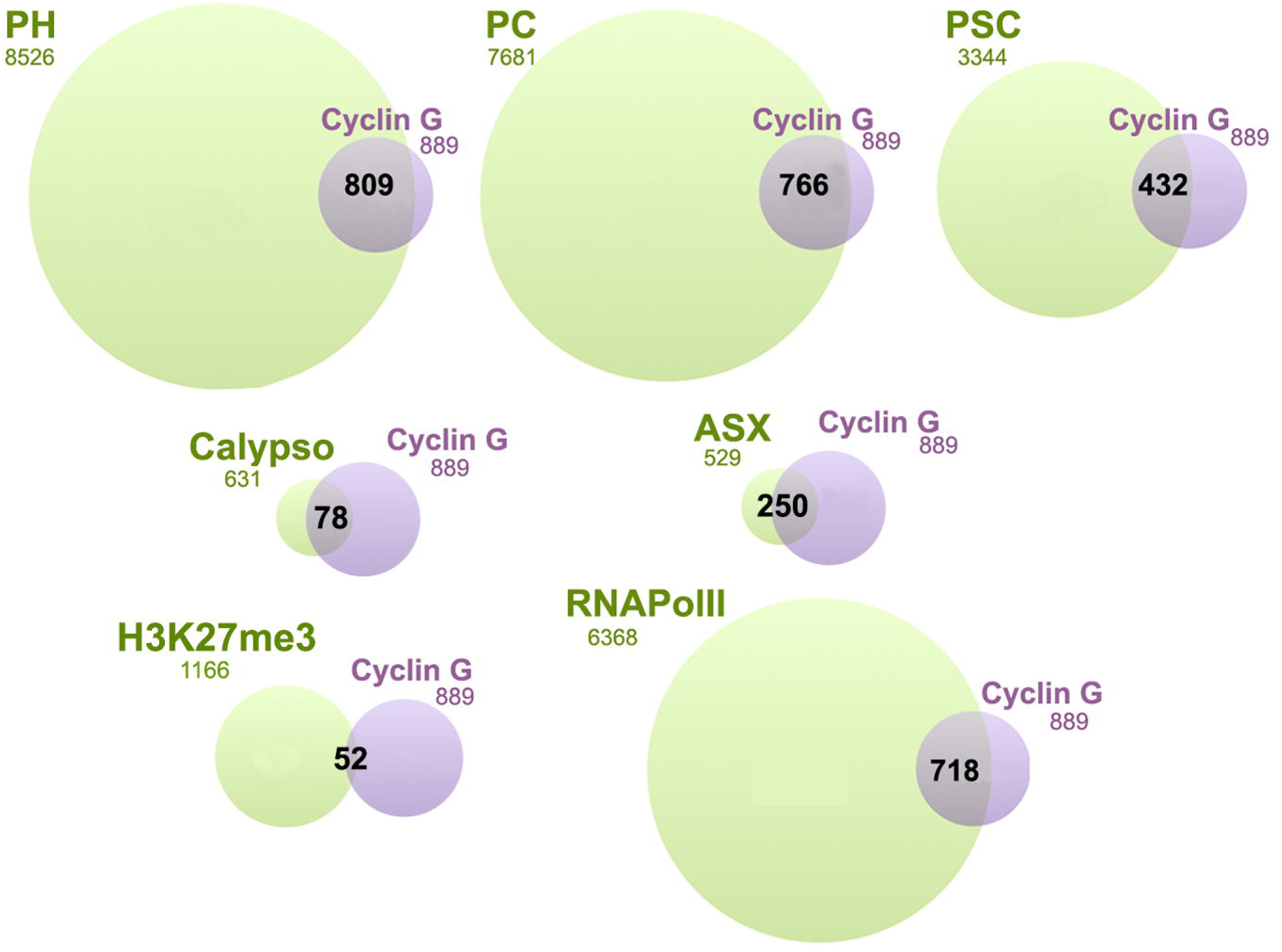
Cyclin G shares target genes with PRC1, ASX and RNAPolII but not with Calypso. Venn diagrams showing the intersection between Cyclin G-bound genes in *+/UAS-CycG*^*ΔP*^; *da-Gal4/+,* wing imaginal discs and PH, PC, PSC, ASX, Calypso and RNApolII in wild-type wing imaginal discs.

### Cyclin G is central in the wing imaginal disc network

These genome-wide analyses indicate that Cyclin G coordinates the expression of genes involved in translation and energy production. However, only a few Cyclin G-bound genes were deregulated in *da-Gal4, UAS-CycG*^*ΔP*^*/+* imaginal discs. To better understand how Cyclin G orchestrates target gene expression, we developed a systems biology approach. We first built an interactome based on genes expressed in control *da-Gal4/+* wing imaginal discs (with a cutoff of 10 reads). Edges corresponding to protein-protein interactions (PPI) and transcription factor-gene interactions (PDI) were integrated into this interactome through DroID (Murali et al. 2011). The resulting wing imaginal disc interactome, further called the WID network, was composed of 9,966 nodes (proteins or genes) connected *via* 56,133 edges (interactions) (WID.xmml). We then examined the position of Cyclin G in this network. Betweenness centrality - *i.e.* the total number of non-redundant shortest paths going through a certain node – is a measure of centrality in a network (Yu et al. 2007). A node with a high betweenness centrality could control the flow of information across the network (Yamada and Bork 2009). With 8.32E-03, Cyclin G had one of the highest value of betweenness centrality of the network, ranking at the 30^th^ position among the 9,966 nodes. This suggests that Cyclin G represents a hub in the WID network.

In order to isolate a connected component of the WID network that showed significant expression change when *CycG*^*ΔP*^ is expressed, we introduced the expression matrix describing expression of the 530 significantly deregulated genes in the WID network. We next used JactiveModules to identify sub-networks of co-deregulated genes (Ideker et al. 2002). A significant sub-network of 222 nodes and 1069 edges centred on Cyclin G was isolated (Z score 48.53). This sub-network was laid out according to functional categories (Fig. 9A, CycG_subnetwork.xmml). Four modules composed of genes respectively involved in transcription, mitochondrial activity, translation, and metabolism, were found to be highly connected to Cyclin G. Strikingly, the “translation” module was mainly composed of genes up-regulated in *da-Gal4, UAS-CycGΔP/+* wing imaginal discs. On the contrary, the “mitochondrion” and “metabolism” modules were mainly composed of genes down-regulated in *da-Gal4, UAS-CycG*^*ΔP*^*/+* wing imaginal discs. This sub-network might be important for developmental stability. Interestingly, Cyclin G-bound genes in this sub-network were enriched in genes bound by the PRC1 proteins PC, PH and PSC, as well as by RNAPolII, and to a lesser extent by ASX (Fig. 9B).

**Figure 9.**
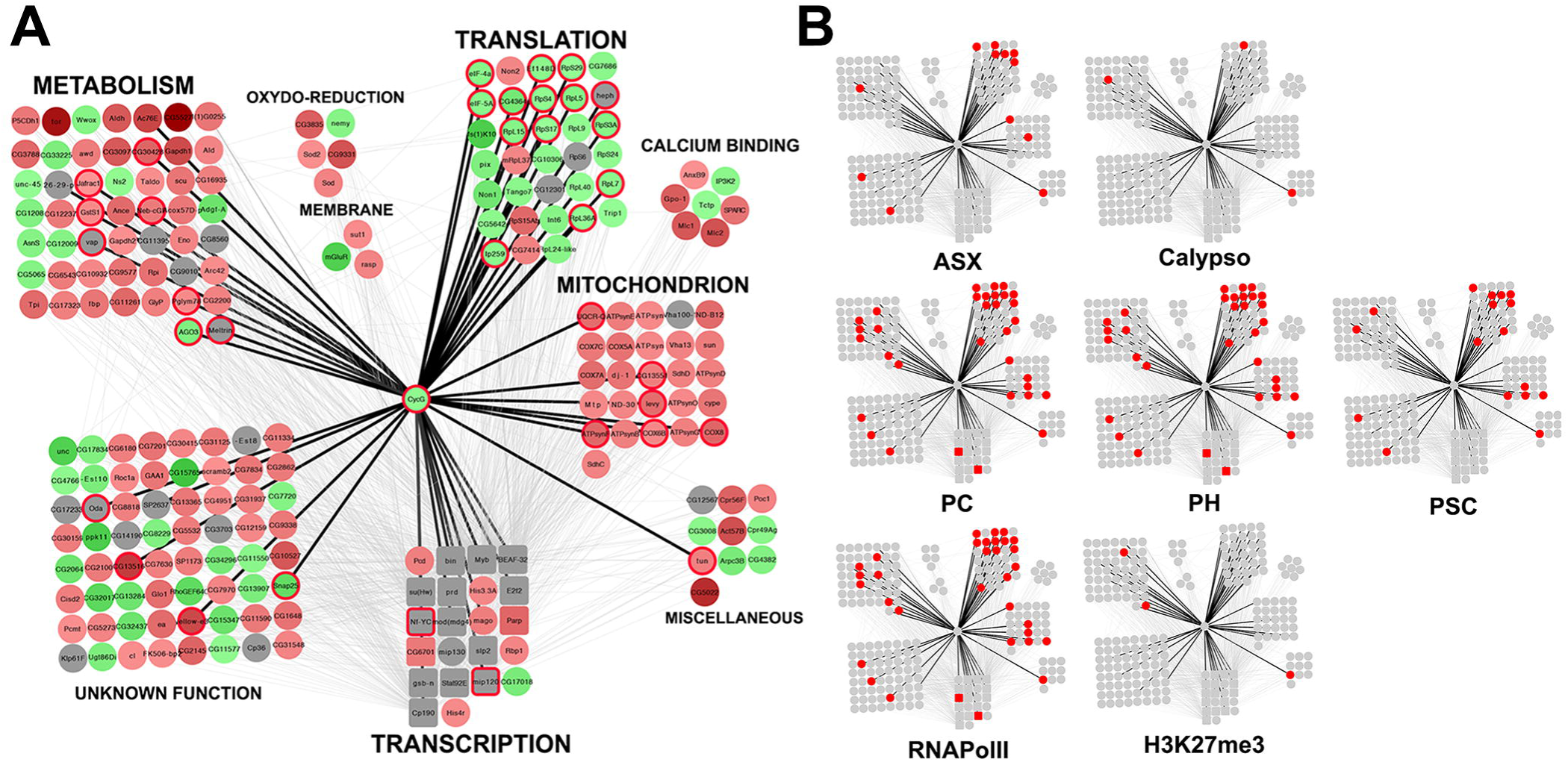
Functional subnetwork identified in wing imaginal discs expressing *CycG^ΔP^*. **A –** Schematic representation of a sub-network of 222 genes centred on Cyclin G (CycG_subnetwork.xmml) and identified using JactiveModules (Z score 48.53). In this sub-network, 65 genes were up-regulated in *da-Gal4, UAS-CycG^ΔP^ vs da-Gal4/+* wing imaginal discs (green gradient), 124 genes were down-regulated (red gradient), and 33 genes were not significantly deregulated (grey). Genes bound by Cyclin G are circled in red. Transcription factor genes are represented by squares. Genes were clustered depending on their function. Black edges correspond to interactions discovered in the present study. Grey edges correspond to interactions described in the literature and imported into the WID network using DroID. **B –** Genes bound by ASX, Calypso, PC, PH, PSC, or RNAPolII, or enriched in H3K27me3 in the sub-network are represented in red.

## Discussion

The *CycG* gene of *D. melanogaster* encodes a cyclin involved in transcriptional control, cell growth and the cell cycle (Salvaing et al. 2008; Faradji et al. 2011). Mild overexpression of a cDNA encoding Cyclin G deleted of a short C-terminal sequence potentially involved in Cyclin G degradation, a PEST-rich domain, induces high FA, notably of wings, providing a unique tool to investigate the genetic bases of developmental stability. (Debat et al. 2011; Debat and Peronnet 2013). Cyclin G interacts physically with two chromatin regulators of the Enhancers of Trithorax and Polycomb family (ETP), and genetically with *Polycomb-group* (*PcG*) genes (Dupont et al. 2015). This prompted us to examine the role of these interactions in developmental stability and to investigate deeply the function of Cyclin G in transcriptional regulation.

### Cyclin G maintains developmental stability through an organ-autonomous process that involves the PRC1 and PR-DUB complexes

In *Drosophila*, mutations of the genes encoding the insulin-like peptide 8 (*Dilp8*) and its receptor (*Lgr3*) that both participate in systemic coordination of growth *via* a well-identified neuronal circuit, have been shown to induce abnormally high FA (Parker and Shingleton 2011; Garelli et al. 2012; Colombani et al. 2012, 2015; Vallejo et al. 2015). To investigate the role of *CycG* in this process, we deregulated it in the different modules of the circuit. *CycG-*induced wing FA only occured when the deregulation was local, *i.e.* in wing imaginal discs. Although we cannot exclude that Cyclin G induces expression of a systemic factor that is dumped into the haemolymph, our observations suggest that *CycG* maintains developmental stability through an autonomous mechanism that would not involve the systemic *Dilp8/Lgr3* pathway. Such a mechanism recalls Garcia-Bellido’s Entelechia model which proposes that local interactions between wing imaginal disc cells orchestrate their own proliferation in order to generate an adult organ of constant size and shape, independently of global cues (García-Bellido and García-Bellido 1998; García-Bellido 2009).

Expression of Cyclin G deleted of the ETP interacting domain doubles FA as compared to expression of Cyclin G with this domain, irrespective of whether the PEST domain is present or not. Hence, the interaction between Cyclin G and chromatin regulators might somehow participate in developmental stability. We show that mutations of the PRC1 and PR-DUB encoding genes strongly increase *CycG-* induced FA. Moreover, many of the genes that are bound by Cyclin G in wing imaginal discs are also bound by PRC1 and by ASX. Altogether, these observations suggest that transcriptional regulation of target genes shared by Cyclin G, PRC1 and ASX is of paramount importance for developmental stability. We did not observe any significant overlap between Cyclin G-bound genes and binding sites for Calypso, the second component of PR-DUB. Yet, *caly* mutations strongly increase *CycG*-induced FA. Thus, the role of PR-DUB in this context remains to be clarified.

PRC1 and PR-DUB contain antagonistic enzymes (SCE/dRing and Calypso) that respectively ubiquitinates and deubiquitinates H2A on lysine 118 in *Drosophila* (lysine 119 in human). Although Cyclin G co-localizes extensively with H2AK118ub on polytene chromosomes, the global level of H2AK118 ubiquitination is not modified in tissues where Cyclin G isoforms are overexpressed suggesting that this epigenetic mark is not involved in developmental stability. It was recently shown that canonical PRC1 accounts for only a small fraction of global H2AK118ub, most of this ubiquitination being due to L(3)73Ah, a homolog of mammalian PCGF3 (Lee et al. 2015). Our data supporting the importance of the interaction between Cyclin G and canonical PRC1, it is tempting to speculate that PRC1 and PR-DUB are partners in the ubiquitination/deubiquitination of an unknown protein important for developmental stability.

### Regulation of growth during the cell cycle might be a factor of developmental stability Drosophila

Cyclin G and the two vertebrate G-type cyclins, CCNG1 and CCNG2, share many features (Salvaing et al. 2008; Fischer et al. 2015, 2016). Furthermore, they exhibit a complex relationship to growth, on the one hand promoting it, (Smith et al. 1997; Kimura et al. 2001; Fischer et al. 2015, 2016) and on the other hand, slowing down or even blocking the cell cycle (Zhao et al. 2003; Arachchige-Don et al. 2006; Faradji et al. 2011; Zimmermann et al. 2016). Strickingly, in *Drosophila,* a *CycG* null mutant (*CycG*^*HR7*^) as well as *CycG* overexpressing flies (*da-Gal4; UAS-CycG* or *da-Gal4; UAS-CycG*^*ΔP*^) exhibit reduced body weight (Debat et al. 2011; Fischer et al. 2015, 2016). However, a very low *CycG* overexpression rescues *CycG*^*HR7*^ body weight (Fischer et al. 2015). In the same vein, whereas reducing *CycG* expression by RNA interference increases shape FA (Debat et al. 2011), we do not observe any increase of size or shape FA in the *CycG*^*HR7*^ null mutant (data not shown).

*CycG*-induced FA is associated with loss of correlation between cell size and cell number, connecting developmental stability to growth regulation during the cell cycle (Debat et al. 2011). As Cyclin G deregulation impairs growth in G1 phase of the cell cycle (Faradji et al., 2011), this supports the idea that a mechanism linked to regulation of cell cycle-dependent growth is essential for developmental stability. The fact that many genes involved in translation, energy production and metabolism are controlled by Cyclin G strengthens this hypothesis.

It was shown that promoters of actively transcribed genes, notably ribosomal protein genes, are bookmarked by ubiquitination during mitosis. This mechanism would allow post-mitotic resumption of their transcription at the very beginning of the G1 phase, thus promoting cell growth. The enzymes responsible for this ubiquitination, whose substrat is unknown but different from H2AK119, are the vertebrate PSC homolog BMI1, and Ring1A, one of the SCE/dRing homologs (Arora et al. 2015). Cyclin G binds the promoter of many ribosomal protein genes and a high FA is observed when Cyclin G lacking the PEST domain, a potential ubiquitination site, is expressed. Hence, an exciting hypothesis would be that Cyclin G is ubiquitinated by PRC1, or PSC and SCE/dRing outside PRC1, and binds the promoter of these genes during mitosis. Its deubiquitination par PR-DUB at the onset of G1 phase would release the transcriptional standby of active genes at the end of mitosis. In agreement with this, we found that genes involved in metabolism and mitochondrial activity are down-regulated in the *CycG*^*ΔP*^ context. However, we observed at the same time that ribosomal protein genes are up-regulated which should rather promote growth. This foreshadows a complex relationship between Cyclin G and the PRC1 and PR-DUB complexes in the cell cycle-dependent regulation of these genes.

### Fine-tuned regulation of genes involved in translation, metabolism and mitochondrial activity is necessary for developmental stability

Cyclin G appears central in a small regulatory sub-network that connects genes involved in metabolism, mitochondrial activity and translation. Besides, many of Cyclin G’s direct transcriptional targets in this network are also targets of PRC1 and RNAPolII, and to a lesser extent of ASX. Interestingly, a large scale analysis of the *Drosophila* wing imaginal disc proteome has recently shown that wing size correlates with some basic metabolic functions, positively with glucose metabolism and negatively with mitochondrial activity, but not with ribosome biogenesis (Okada et al. 2016). In agreement with this, we report here that many genes involved in basic metabolism, such as for example *Gapdh1, Gapdh2* or *Jafrac1,* are down-regulated in the *CycGΔP* context, which also agrees with the small mean size of *CycGΔP* flies, organs and cells. However, while mitochondrial genes are negatively regulated, ribosomal biogenesis genes are simultaneously positively regulated. Although transcriptome variations are probably not a direct image of proteome variations, our data suggest that robustness of wing size correlates with the fine-tuning of these key functions relative to each other.

### Noisiness of gene expression as a source of developmental noise

Cyclin G, PRC1 and PR-DUB are mainly involved in the regulation of transcription. *CycG*-induced high FA is associated with high variability of cell size, that might be due to variability in expression of target genes that are mainly involved in growth control. An exciting hypothesis would be that alteration of developmental stability is due to the noisy transcription of their shared targets. Phenotypic variations in isogenic populations of both prokaryotic and eukaryotic cells may indeed result from stochastic gene expression mechanisms (McAdams and Arkin 1997). An increasing corpus of data suggests that the process of gene regulation *per se* can strongly affect variability in gene expression among adjacent cells (for a review see Sanchez et al. 2013). Interestingly, in several cases, noise in gene expression specifically concerns a subset of genes (Weinberger et al. 2012). Transcriptional noise may arise at all steps of transcription. For example, RNA polymerase II pausing during elongation is a source of transcriptional noise (Rajala et al. 2010). However, the binding of Cyclin G on many TSS is rather in favor of a role in limiting noisy initiation of transcription. Recently, activity of the Polycomb complex PRC2 was shown to be important to prevent spurious transcription of inactive genes and suppress pervasive transcription of intergenic regions (Lee et al. 2015). Mutations of *E(z)* and *esc* that encode two PRC2 members has no effect on *CycG*-induced FA. Dysfunction of PRC2-dependent spurious transcription control is thus unlikely to be the cause of any *CycG*-induced developmental noise. Nevertheless, a similar but weaker effect on intergenic transcription was attributed to PRC1 (Lee et al. 2015). It is thus tempting to speculate that cooperation between Cyclin G and the PRC1 and PR-DUB complexes is important to prevent spurious transcription of genes involved in growth in the broad sense. It will be very interesting to address these points in the future.

## Material and Methods

### Plasmids

The *pPMW-attB* plasmid was built as follows: the Gateway® vector *pPMW* (Huynh and Zieler 1999) was linearized by digestion with *NsiI*; the *attB* sequence was amplified from *pUASTattB* (Bischof et al. 2007) using primers *attB-NsiIF* and *attB-NsiIR* (Sup. Table 15) and the PCR product was digested with *NsiI*; the digested PCR product and the linearized plasmid were ligated and sequenced. This plasmid was deposited at Addgene (plasmid # 61814).

The full-length *CycG* cDNA (*CycG*^*FL*^, encoding the 566 amino-acid protein) was amplified from S2 cell cDNAs using primers *CycGnF* and *CycGnR*. cDNAs encoding truncated forms of Cyclin G (*CycG*^*ΔP*^, Cyclin G deleted of the putative PEST domain corresponding to amino-acids 542 to 566; *CycG*^*ΔE*^, Cyclin G deleted of the ETP-interacting domain corresponding to amino-acids 1 to 130; *CycG^ΔEΔP^*, Cyclin G deleted of both domains) were amplified from the full-length *CycG* cDNA using primers *CycGnF* and *CycG541R*, *CycG130F* and *CycGnR*, and *CycG130F* and *CycG541R*, respectively (Sup. Table 15). The PCR products were cloned into *pENTR/D-TOPO^®^* (Invitrogen), transferred into *pPMW-attB* and the resulting plasmids *pPMW-attB-CycG^FL^*, *pPMW-attB-CycG^ΔP^*, *pPMW-attB-CycG^ΔE^*, *pPMW-attB-CycG^ΔEΔP^* were sequenced.

### Drosophila melanogaster strains and genetics

Flies were raised on standard yeast-cornmeal medium at 25°C.

*UAS-Myc-CycG* transgenic lines were obtained by *PhiC31*-integrase mediated insertion into strain *y*^*1*^*M{vas-int.Dm}ZH-2Aw*^*^;*M{3xP3-RFP.attP’}ZH-51C* (stock BL-24482). Plasmids *pPMW-attB-CycG^FL^*, *pPMW-attB-CycG*^*ΔP*^, *pPMW-attB-CycG*^*ΔE*^ and *pPMW-attB-CycG*^*ΔEΔP*^ were injected into embryos, G0 adults were back-crossed to *yw*, and G1 transformants were crossed to *yw* again to obtain G2 transformants (BestGene Inc.). Transformants were individually crossed with *yw; Sp/CyO*, and the curly wing siblings were crossed with each other. Homozygous transgenic lines were then obtained by crossing 5 females and 5 males. The resulting lines were named *UAS-CycG^FL^, UAS-CycG*^*ΔP*^, *UAS-CycG*^*ΔE*^ and *UAS-CycG*^*ΔEΔP*^.

Gal4 drivers used were *daughterless-Gal4* (*da-Gal4*), *nubbin-Gal4* (*nub-Gal4*), *optomotor-blind-Gal4* (*omb-Gal4*), *rotund-Gal4* (*rn-Gal4*), *scalloped-Gal4* (*sd-Gal4*), *teashirt-Gal4* (*tsh-Gal4*), *vestigial-Gal4* (*vg-Gal4*) (from the Bloomington *Drosophila* stock center), and *Insulin-like peptide 3-Gal4* (*dILP3-Gal4*), *neuropeptide F-Gal4* (*NPF-Gal4), Pigment-dispersing factor-Gal4* (*Pdf-Gal4*), *period-Gal4* (*per-Gal4*), *phantom-Gal4* (*phm-Gal4*), *Prothoracicotropic hormone-Gal4* (*Ptth-Gal4*), *R10B09-Gal4* (kind gifts from Dr M. Dominguez’s lab).

The *da-Gal4, UAS-CycG^ΔP^* third chromosome, obtained by recombination of *da-Gal4* with the original *UAS-CycG*^*ΔP*^ transgene (*RCG76*), was used to test genetic interactions between *CycG* and several *PcG* or *ETP* mutations (Dupont et al. 2015). *PcG* and *ETP* alleles used are described in Table 1.

For FA analyses, five replicate crosses were performed for each genotype, wherein 6 females carrying a Gal4 driver were mated with 5 males carrying a *CycG* transgene. Parents were transferred into a new vial every 48 h (three times) then discarded. Thirty females were sampled from the total offspring of the desired genotype. For genetic interactions with *PcG* or *ETP* mutants, crosses were performed similarly except that 6 *PcG* or *ETP* mutant females were mated either with 5 *da-Gal4, UAS-CycG*^*ΔP*^ males, or with 5 *da-Gal4* males as control.

### Morphometrics

Right and left wings of 30 sampled females were mounted on slides, dorsal side up, in Hoyer’s medium. Slides were scanned with a Hamamatsu Nanozoomer Digital Slide scanner. Wing pictures were exported into tif format using NDP.view. All wings were oriented with the hinge to the left. Image J was used to digitize 15 landmarks or only landmarks 3 and 13 when indicated (Fig. 1B). All wings were measured twice. Analysis of wing size FA, the variance of the difference between the left and the right wing in a population, was performed as described previously using the Rmorph package (Debat et al. 2011). The FA10 index was used to estimate FA, *i.e.* FA corrected for measurement error, directional asymmetry and inter-individual variation (Palmer and Strobeck 1992). For all genotypes, the interaction individual*side was significant, indicating that FA was larger than measurement error. F-tests were performed to compare the different genotypes.

### Immunostaining of polytene chomosomes and wing imaginal discs

Immunostainings were performed as described in Dupont et al. (2015). Primary anti-H2AK119ub antibodies (Cell Signaling D27C4) were used at a 1:40 dilution.

### RNA-seq experiments and RT-qPCR validations

Wing imaginal discs from *da-Gal4/UAS-CycG^ΔP^* and *da-Gal4/+* third instar female larvae were dissected, and total RNAs were extracted as previously described except that 150 discs homogenized by pipetting were used for each extraction (Coléno-Costes et al. 2012). Three biological replicates (wing imaginal discs dissected from three independent crosses) were generated for each genotype. Library preparation and Illumina sequencing were performed at the ENS Genomic Platform (Paris, France). PolyA RNAs were purified from 1 μg of total RNA using oligo(dT). Libraries were prepared using the TruSeq Stranded mRNA kit (Illumina). Libraries were multiplexed by 6 on 2 flowcell lanes. 50 bp single read sequencing was performed on a HiSeq 1500 device (Illumina). Number of reads are shown on Sup. Table 16. Reads were aligned with the *D. melanogaster* genome (dm6, r6.07) using TopHat 2 (v2.0.10) (Kim et al. 2013). Unambiguously mapping reads were then assigned to genes and exons described by the Ensembl BDGP5 v77 assembly, by using the “summarizeOverlaps” function from the “GenomicAlignments” package (v 1.2.2) in “Union” mode (Lawrence et al. 2013). Library size normalization and differential expression analysis were both performed with DESeq 2 (v 1.6.3). Genes with adjusted p-value below 0.05 were retained as differentially expressed (Love et al. 2014). Gene Ontology analysis was performed using DAVID (Huang et al., 2009).

For RT-qPCR validations, RNAs were extracted from wing imaginal discs and treated with Turbo DNAse (Ambion). cDNA were synthesized with SuperScript II Reverse transcriptase (Invitrogen) using random primers. RT-qPCR experiments were carried out in a CFX96 system (Bio-Rad) using SsoFast EvaGreen Supermix (Bio-Rad). Two biological replicates (cDNA from wing imaginal discs of larvae coming from independent crosses) and three technical replicates (same pool of cDNA) per biological replicate were performed for each genotype. Expression levels were quantified with the Pfaffl method (Bustin et al. 2009). The geometric mean of two reference genes, *Lamin* (*Lam*) and *rasputin* (*rin*), the expression of which did not vary when *CycG*^*ΔP*^ was expressed, was used for normalization (Vandesompele et al. 2002). Sequences of primer couples are listed in Sup. Table 15.

An interactome was built using Cytoscape (v 2.8.3) and the DroID plugin (v 1.5) to introduce protein-protein and transcription factor-gene interactions (Murali et al. 2011). The jActiveModules plugin (v 2.23) was used to find sub-networks of co-deregulated genes in the interactome by using “overlap threshold” 0.8, “score adjusted for size”, and “regional scoring” (Ideker et al. 2002).

### ChIP-seq experiments and ChIP-qPCR validations

Wing imaginal discs from *+/UAS-CycG*^*ΔP*^; *+/da-Gal4* and *+/da-Gal4* third instar female larvae were used for ChIP-seq experiments. 600 wing imaginal discs were dissected (taking one disc per larva) in Schneider medium and aliquoted per 50 on ice. The 12 microtubes were treated as described in (Coléno-Costes et al. 2012) with minor modifications. Discs were fixed at 22°C. 12 sonication cycles were performed (Diagenode Bioruptor sonifier; cycles of 30’’ ON, 30’’ OFF, high power). After centrifugation, the 12 supernatants were pooled, homogenized, and 2% were kept (Input). The remaining fragmented chromatin was redistributed into 12 tubes and each tube was adjusted to 1 mL with 140 mM NaCl, 10 mM Tris-HCl pH 8.0, 1 mM EDTA, 1% Triton X-100, 0.1% sodium deoxycholate, 0.1% BSA, Roche complete EDTA-free protease inhibitor cocktail. For immunoprecipitation, 3 μg of anti-Myc antibody (Abcam 9132) were added per tube. Two biological replicates were performed.

Library preparation and Illumina sequencing were performed at the ENS Genomic Platform (Paris, France). Libraries were prepared using NEXTflex ChIP-Seq Kit (Bioo Scientific), using 38 ng of IP or Input DNA. Libraries were multiplexed by 10 on one flowcell run. 75 bp single read sequencing was performed on a NextSeq 500 device (Illumina). Reads were filtered by the “fastq_quality_filter” command from the “fastx-Toolkit” package (http://hannonlab.cshl.edu/fastx_toolkit/), using a threshold of 90% bases with mapping quality ≥ 20. Reads are shown on Sup. Table 17. Those that successfully passed the filtering step were aligned to the *D. melanogaster* genome (dm6, r6.07) using Bowtie 2 (http://bowtie-bio.sourceforge.net/bowtie2/) (v 2.1.0) with default parameters (Langmead and Salzberg 2012). Peaks were called by MACS2 (v 2.1.0) by comparing each ChIP to its input library, with fragment size fixed at 110 bp and otherwise default parameters (Zhang et al. 2008). Peak reproducibility between the two biological replicates was then analysed with the IDR method (https://www.encodeproject.org/software/idr/) (Li et al. 2011). Briefly, an IDR score was assigned to each peak by the “batch-consistency-analysis” function, using the recommended parameters for MACS peaks (peak ranking based on p-value). Peaks below the 0.05 threshold were considered reproducible. The overlapping reproducible peaks from both replicates were fused using the BEDtools suite “merge” function (Quinlan and Hall 2010), resulting in the final list of peaks kept for subsequent analysis. Cyclin G-bound genes were defined as genes from the genome annotation file (dm6, r6.07) which overlapped at least one of these Cyclin G peaks, as obtained by the BEDtools suite “intersect” function.

For ChIP-qPCR validations, ChIP were performed similarly with the anti-Myc antibody. Rabbit IgG (Diagenode) were used as negative control (mock). qPCR experiments were carried out in a CFX96 system (Bio-Rad) using SsoFast EvaGreen Supermix (Bio-Rad). Three biological replicates – three technical replicates per biological replicate – were performed for each antibody and for the Input. Sequences of primer couples are listed in Sup. Table 15. Data were normalized against Input chromatin.

Heatmaps and aggregation plots of Cyclin G signal over gene bodies and TSS were generated using the ngsplot package. (https://github.com/shenlab-sinai/ngsplot) (Shen et al. 2014). Some genes with spurious signal (such as genes from the histone complex) were excluded from the analysis based on signal uniformity over the full length of the gene (cumulative derivative of Cyclin G signal over gene length = 0).

## Data access

Gene Expression Omnibus accession numbers for RNA-seq data: GSE99462, GSM2644389, GSM2644390, GSM2644391, GSM2644392, GSM2644393, GSM2644394; for ChIP-seq data: GSE99461, GSM2644385, GSM2644386, GSM2644387, GSM2644388.

### Genomic association

Genomic loci enriched for Polycomb (PC), Posterior Sex Comb (PSC), Polyhomeotic (PH), RNA Polymerase II (RNAPolII) and H3K27me3 in wild type imaginal discs of third instar larvae were retrieved from GEO (GSE42106) (Schaaf et al. 2013) (H3K27me3_WholeWingDisc GSM1032567, PcRJ_AnteriorWingDisc GSM1032571, PcRJ_PosteriorWingDisc GSM1032574, Ph_WholeWingDisc GSM1032576, PolII_WholeWingDisc GSM1032577, Psc_WholeWingDisc GSM1032578. Binding sites for PC in the whole wing disc were defined as the overlap between PC binding sites in the anterior and posterior wing disc compartment, as obtained by the BEDtools “intersect” function. For ASX and Calypso, the bed files were a kind gift from Dr. J. Müller (Scheuermann et al. 2010). The mappability file for dm6 genome with 25 nt reads (the smallest size in the compared data) was generated using the Peakseq code (http://archive.gersteinlab.org/proj/PeakSeq/Mappability_Map/Code). The overall size of the mappable genome was used as the effective genome size for the GAT software (https://github.com/AndreasHeger/gat) to assess the significance of the overlap between peaks of Cyclin G and other factors (Heger et al. 2013). As GAT performs a two-tailed test, it displays low p-values both for significant overlap and exclusion (as between Cyclin G and H3K27me3).

Gene overlap significance assessment was made as follows: under the null hypothesis, genes that are enriched for ASX, Calypso, PC, PSC, PH, RNAPolII or H3K27me3 in wild type imaginal discs of third instar larvae should not exhibit any bias towards Cyclin G targets. Thus, the overlap between n enriched genes and K Cyclin G targets genes should be explained by random sampling without replacement of n genes within the total amount N of *D. melanogaster* genes. The amount of overlap under the null hypothesis X follows a hypergeometric law: *X∼HY(K, N, n)*. The significance of the observed overlap k was computed as the probability of observing X higher or equal to k under the null hypothesis: P(X ≥ k).

## Acknowledgments

We thank Dr. E. Mouchel-Vielh and Dr. J-M. Gibert for stimulating discussions and critical reading of the manuscript, the Bloomington Stock Center for fly strains, Dr. J. Müller for the alleles of *Asx* and *caly* and for the ASX and Calypso ChIP bed files, Dr. M. Dominguez and Dr S. Juarez-Carreño for Gal4 drivers. This work was funded by the Centre National de la Recherche Scientifique (CNRS), Université Pierre et Marie Curie (UPMC), Sorbonne Universités (grant SU-14-R-CDV-05-1 to FP), and Fondation ARC pour la Recherche sur le Cancer (grant PJA20131200314 to FP). The école Normale Supérieure genomic platform was supported by the France Génomique national infrastructure, funded as part of the “Investissements d’Avenir” program managed by the Agence Nationale de la Recherche (contract ANR-10-INBS-09). CAD was funded by a doctoral fellowship from the MESR (Ministère de l’Enseignement Supérieur et de la Recherche). JD was funded by a doctoral fellowship from the MESR and by the Fondation pour la Recherche médicale (FDT20160435164).

